# Intra-organ cell specific mitochondrial quantitative interactomics

**DOI:** 10.1101/2024.06.10.598354

**Authors:** AA Bakhtina, MD Campbell, BD Sibley, M Sanchez-Contreras, MT Sweetwyne, JE Bruce

## Abstract

Almost every organ consists of many cell types, each with its unique functions. Proteomes of these cell types are thus unique too. But it is reasonable to assume that interactome (inter and intra molecular interactions of proteins) are also distinct since protein interactions are what ultimately carry out the function. Podocytes and tubules are two cell types within kidney with vastly different functions: podocytes envelop the blood vessels in the glomerulus and act as filters while tubules are located downstream of the glomerulus and are responsible for reabsorption of important nutrients. It has been long known that for tubules mitochondria plays an important role as they require a lot of energy to carry out their functions. In podocytes, however, it has been assumed that mitochondria might not matter as much in both normal physiology and pathology^1^. Here we have applied quantitative cross-linking mass spectrometry to compare mitochondrial interactomes of tubules and podocytes using a transgenic mitochondrial tagging strategy to immunoprecipitate cell-specific mitochondria directly from whole kidney. We have uncovered that mitochondrial proteomes of these cell types are quite similar, although still showing unique features that correspond to known functions, such as high energy production through TCA cycle in tubules and requirements for detoxification in podocytes which are on the frontline of filtration where they encounter toxic compounds and therefore, as a non-renewing cell type they must have ways to protect themselves from cellular toxicity. But we gained much deeper insight with the interactomics data. We were able to find pathways differentially regulated in podocytes and tubules based on changing cross-link levels and not just protein levels. Among these pathways are betaine metabolism, lysine degradation, and many others. We have also demonstrated how quantitative interactomics could be used to detect different activity levels of an enzyme even when protein abundances of it are the same between cell types. We have validated this finding with an orthogonal activity assay. Overall, this work presents a new view of mitochondrial biology for two important, but functionally distinct, cell types within the mouse kidney showing both similarities and unique features. This data can continue to be explored to find new aspects of mitochondrial biology, especially in podocytes, where mitochondria has been understudied. In the future this methodology can also be applied to other organs to uncover differences in the function of cell types.

## INTRODUCTION

Over the last few years omics fields such as genomics, transcriptomics and proteomics have been moving toward studying cell type heterogeneity. This was made possible by great advances in single cell technologies.^2,3^ Interactomics, or study of intra and inter protein interactions on a system level usually by the means of cross-linking mass spectrometry (XL-MS), has inherent constraints that make move toward single cell studies particularly challenging. Cross-links are formed by addition of a cross-linking reagent reactive toward specific protein residues (most commonly lysines), with low efficiency and require high input amounts to be detected^4^. Although, cell typed specific protein-protein interactions in cultured cell lines have been studied by other means, such AP-MS^5^, there are no reported cell type specific interactomes stemming from the same animal organ. Making interactome data quantitative (i.e. finding differences that are not just present/absent but have a more subtle variation) is an even bigger challenge^6^. Here, we report the first quantitative comparison of mitochondrial interactomes of two cell types originating in the same organ substructure: podocytes and tubules from kidney nephrons. By combining quantitative XL-MS (qXL-MS) technique that utilizes isobaric quantitative Protein Interaction Reporter (iqPIR)^7^ and novel mouse models that express hemagglutinin (HA) tag on mitochondrial outer membrane of kidney cells (unpublished data) in cell type specific manner, we were able to detect quantitative differences in the interactomes of podocytes and tubules. qXL-MS allows for the detection of changes that are interaction (protein-protein and protein-ligand), conformation, and lysine post-translational modification (PTM) driven. Our method showed remarkable reproducibility between biological replicates with the lowest R^2^ value of 0.8. By adding DIA-based quantitative proteome comparisons we established interactome differences that are independent of protein abundances.

Mitochondrial proteome differences between podocytes and tubules offered some insights but pathway analysis was limited as only 45 mitochondrial proteins are differentially expressed. On the other hand, utilizing proteins with differential cross-link abundances allowed us to perform pathway enrichment analysis and we were able to identify several pathways that were differentially regulated including betaine metabolism, lysine metabolism, and tryptophan metabolism. This shows that mitochondria play an important albeit distinct role in podocytes compared to tubules.

We have also compared abundances of all, and not just mitochondrial proteins, between podocyte and tubule pull-downs. Lysosomal, cytoskeletal, and general cytosolic proteins were enriched in podocyte pull-downs, while almost all proteins showing higher abundance in tubule pulldowns were mitochondrial. This indicates that podocyte mitochondria might have more contacts with other cellular structures compared to tubule mitochondria.

Previously reported differences in cross-links in antenna region of glutamate dehydrogenase (DHE3) and their relation to the activity^8^ as predicted by protein structure were detected between podocytes and tubules. Predicted differences in DHE3 activity between podocytes and tubules were confirmed orthogonally with colorimetric activity assay.

Overall, this work demonstrates the ability to quantitatively compare interactomes of mitochondria originating from different cell types within an organ. Moreover, insights gained into underexplored mitochondrial biology of podocytes open the door to further studies that can elucidate role of mitochondria in podocyte physiology and podocyte dependent kidney pathology.

## RESULTS AND DISCUSSION

### Quantification of interactome differences between podocyte and tubule mitochondria

Mitochondria was isolated from mouse kidneys with HA tag on either podocyte mitochondria (NPHS2-mtHA) or tubule mitochondria (CDH16-mtHA) using affinity purification column. Mitochondria from podocytes was cross-linked with reporter heavy (RH) cross-linker and tubule mitochondria was cross-linked with stump heavy (SH) cross-linker. This cross-linking scheme allows for comparison of one podocyte sample to one tubule sample. A podocyte mitochondria isolation from a single mouse was combined with a tubule mitochondria isolation from a single mouse, creating three pairs (P1 = pod1/tub1, P2 = pod2/tub2, P3 = pod3/tub3) in 1:1 ratio. Each combined sample was processed and analyzed with mass spectrometry. To identify mitochondrial cross-linked peptides (also referred as cross-links), the data was searched against mitochondrial database Mitocarta 3.0^9^. The iqPIR algorithm was applied to calculate log_2_ ratios for all cross-links detected. In the iqPIR algorithm every peptide and peptide fragment ion peak hat bears a cross-linker stump can be used for quantitation, increasing robustness of the quantitation^7^. This allowed for relative quantitation of cross-links between podocyte and tubule samples in each pair **Figure 1A**). We were able to identify 5119 cross-links and confidently (at least 4 ions contributing to the ratio and 95% confidence interval of the log_2_ ratio distributions < 0.5) quantify 835, 1994 and 1996 in P1, P2 and P3 respectively (**Figure 1B, Supp. Figure 1A**). We then calculated a combined ratio based on ions from all replicates. Filtering for combined ratios that also have at least 2 individual ratios resulted in 2040 total ratios. Pair P1 had lower starting protein amount than pairs P2 and P3. Due to this, and sample processing issues, P1 had fewer identified and quantified cross-links than P2 and P3. Log_2_ ratios for each pair are approximately normally distributed with the majority of cross-links not changing as expected (**Supp. Figure 1B**). We were able to achieve great reproducibility between the pairs as indicated by R^2^ values of 0.80, 0.82 and 0.84 (**Figure 1C**). Cross-links that are decreased, increased or do not change have been quantified as shown in a heatmap of cross-links quantified in at least two out of three pairs (**Figure 1D**). Applying Student’s t-test with Bonferroni correction, we were able to identify cross-links with ratios different from 0 (|log_2_ ratio| ≥0.5) with statistical significance (adj. p-value ≤0.01) (**Figure 1E**).

**Figure 1.**
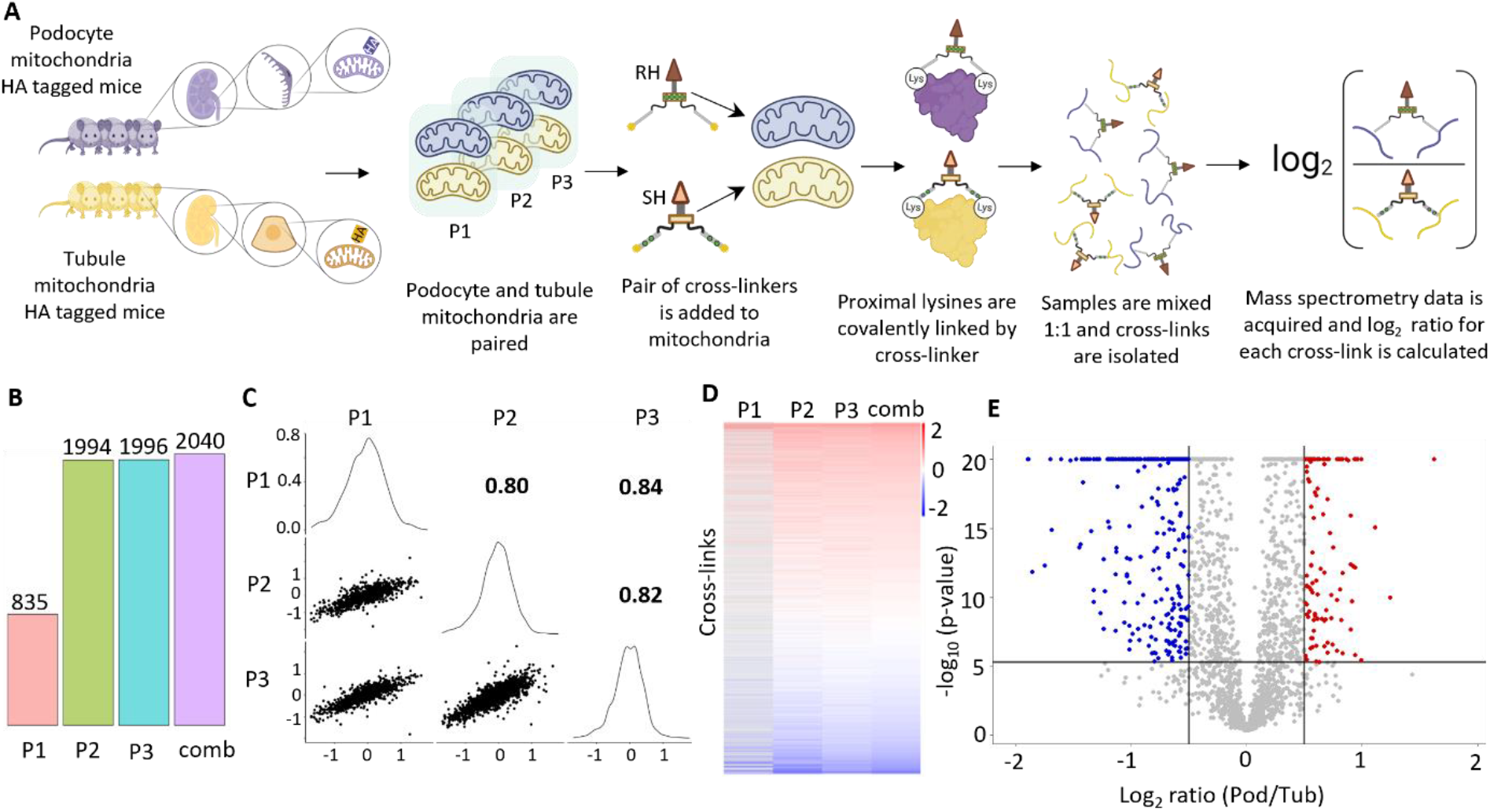
Quantitative comparison of tubules and podocytes interactomes. **A**. Quantitative cross-linking with iqPIR workflow. Mitochondria were isolated from mice expressing the HA tag on the outer mitochondrial membrane in either podocytes or tubules. Mitochondria from podocytes was then cross-linked with reporter heavy (RH) iqPIR reagent and from tubules with stump heavy (SH) reagent to produce a ratio for each cross-link. **B**. Number of confidently quantified cross-links (95% confidence ≤0.5 and at least 4 ions used for quantitation) in each pair (P1-P3) and number of cross-links with confident combined ratio (comb, at least 2 out of three individual confident ratios). **C**. Pairwise correlations between confidently quantified cross-links with Pearson’s R^2^ values indicated. **D**. Heatmap of confidently quantified cross-links shows great agreement among the pairs. **E**. Volcano plot of cross-links log_2_ ratios with 0.05 Bonferroni corrected p-value from one-sample t-test and 0.5 log_2_ ratio used to determine significance.

To complement quantitative interactome data we also acquired proteome measurements for the same samples utilizing DIA-based quantitative proteomics workflow^10,11^. Having information about protein level changes allowed us to distinguish cross-links that are different due to changes in protein abundance from the cross-links that are different due to protein-protein interactions, conformations, protein-ligand interactions or PTMs. Utilizing chromatographic libraries and DIA based quantitation we have identified and quantified 779 protein groups searching against mitochondrial proteome database Mitocarta^9^ (**Supp. Figure 1C**,**D**). Overall, mitochondrial proteomes of tubules and podocytes are fairly similar with only 45 proteins out of 779 being differentially expressed (Benjamin-Hochberg corrected p-value ≤0.05 and |FC| ≥0.5, **Supp. Figure 1E)**. This result is somewhat surprising considering that generally tubules are believed to have greater reliance on mitochondria than podocytes^1^. This could partially be explained by the fact that we normalize mitochondrial protein amount, and it is known that mitochondrial density is lower in podocytes^12^. So, even if podocyte mitochondrial content is lower, the mitochondrial proteomes look similar. Overall, with only 21 proteins significantly up in podocytes and 24 in tubules, it is challenging to perform a meaningful pathway enrichment analysis, but tubule mitochondria show upregulation of TCA cycle and respiratory chain complex I assembly, while cellular detoxification and antioxidant activity is upregulated in podocytes (**Supp. Figure 1F**). While the number of differential proteins is low for an extensive pathway analysis, these results generally agree with functions of tubules and podocytes. Tubules have high energy demand due to active transport across the cell membrane. Podocytes are located at the interface with blood to be filtered by the kidney, so a robust antioxidant system that can help counteract cytotoxicity is needed.

Genetically encoded affinity tags, such as HA, placed at outer mitochondrial membrane, allow for enrichment of the organelle from the tissue homogenate, but this enrichment is not perfect and organelles and proteins that surround the mitochondria are also pulled down. To investigate how this background compares between podocyte and tubule mitochondria we have searched both interactomics and proteomics data against the whole mouse database. We have quantified 2024 proteins with 943 proteins being differentially expressed (Error! Reference source not found. **Figure 2A**), a much larger number than in our mitochondrial searches. We have performed pathway enrichment analysis that showed that the majority of proteins (95%) increased in tubule mitochondria are mitochondrial (**Supp. Figure 2B**), while proteins up in podocyte mitochondria are lysosomal (28%), cytosolic (79%) and cytoskeletal (18%) (Error! Reference source not found. **Figure 2C**). The upregulation of mitochondrial proteins and pathways in tubules thus reflects higher mitochondrial content in the pull-down from tubules and not differential expression of proteins in the mitochondria. At the same time, mitochondrial pull-down in podocytes has a large amount of non-mitochondrial proteins. It is particularly interesting that lysosomal proteins represent a large portion of the non-mitochondrial proteins in podocyte pull-downs. This is in agreement with previous work (unpublished data), where western blots were used to compare abundance of lysosomal marker LAMP1 between the pull-downs. Quantitative proteomics done in this study also shows higher abundance of LAMP1 in podocyte pull-downs (**Supp. Figure 2D)**. Similarly, components of the lysosomal proton pump, V-ATPase, also have higher abundance in podocyte pull-downs (**Supp. Figure 2D**). On a cross-link level we see a similar increase in lysosomal proton pump cross-links (**Supp. Figure 3**). Besides intra-links in lysosomal pump subunits VATA, VATE1, VATB2, we have also detected and quantified inter-links connecting the pump subunits, indicating that the intact lysosomal proton pump has been pulled down and not just individual subunits. Taken together our data show that podocyte mitochondria might have more lysosomal contacts as indicated by an increase in lysosomal protein abundances and cross-links. Lysosomal clearance of damaged mitochondria is important for podocytes since these are terminally differentiated cells that are under constant cytotoxic stress, so abundance of lysosomes near mitochondria could be expected.

**Figure 2.**
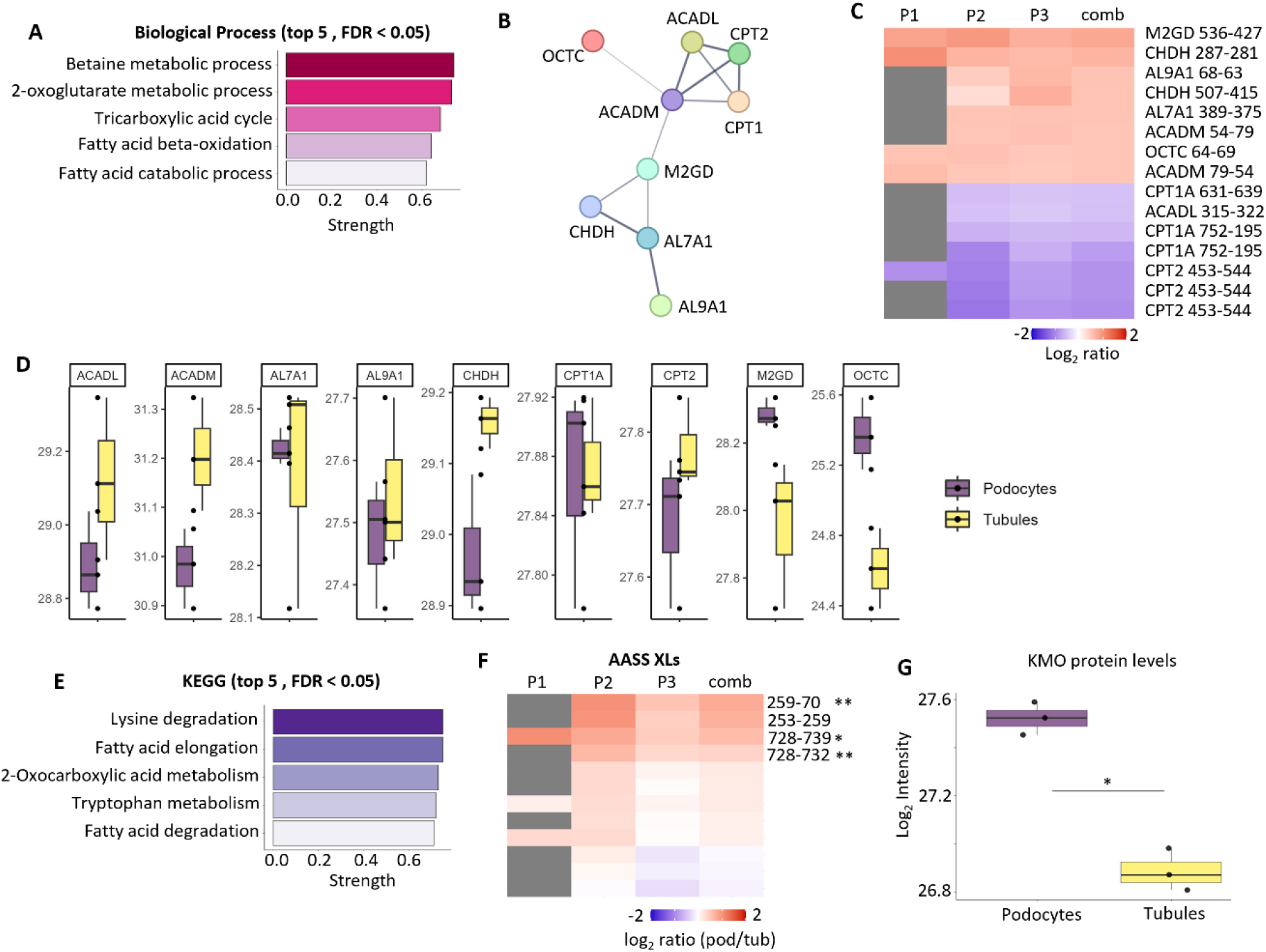
Pathway analysis of differential cross-links. **A**. Top 5 biological processes differential in podocytes and tubules. **B**. Proteins in betaine metabolic pathway that have differential cross-links. **C**. Heatmap of differential cross-links from betaine metabolic pathway. **D**. Protein level comparison of betaine metabolic process proteins that had differential cross-links. **E**. Top 5 KEGG pathways identified. **F**. Heatmap of all cross-links in AASS from lysine degradation and tryptophan metabolism pathways. **G**. Boxplot showing protein levels of KMO in podocytes and tubules. Significance is determined by a two-sided t-test with Benjamin-Hochberg correction at p-value = 0.05.

### Significantly changed cross-links identify mitochondrial pathways different between podocytes and tubules

While mitochondrial proteomes do not show a large number of proteins changing, the main advantage of this study is quantitative interactome data that allows us to leverage knowledge of differences in protein interactions and conformations even when protein levels do not change. We used proteins with significantly changed cross-links (both increased and decreased in podocytes compared to tubules) to perform a pathway enrichment analysis. The top gene ontology biological process was identified as betaine metabolic process (**Figure2A**). Cross-links belonging to 9 proteins (**Figure2B**) from this pathway have been identified as significantly changed with both increased and decreased cross-links present (**Figure 2C**). None of the proteins in this pathway show significantly changed abundancies (OCTC, peroxisomal carnitine O-octanoyltransferase, has the biggest change with log_2_ FC = 0.76 and adjusted p-value = 0.07), so this pathway wouldn’t have been identified as significant based on proteome data alone (**Figure 2D**). Betaine (also known as trimethylglycine) is an amino acid that is important for osmotic regulation^13^. In kidney, it has been shown that supplementation with betaine can have a protective effect during chronic kidney disease and doxorubicin induced toxicity^14,15^. Investigating further the differences in betaine processing between podocytes and tubules could shed light on mechanisms of betaine protection and its connection to podocytes.

Another pathway that we could not have identified without interactomics data is lysine degradation, which is the top hit in KEGG (Kyoto Encyclopedia of Genes and Genomes) analysis (**Figure 2E, Supp. Figure 4A-C**). Kidney plays an integral role in lysine metabolism^16^. The role of lysine in chronic kidney disease has also been documented^17^. Effect of lysine supplementation on a kidney proteome has been investigated and one of the enzymes highlighted is AASS, which catalyzes the first step in lysine degradation through ε-deamination^16^. AASS is composed of two domains: lysine-2-oxoglutarate reductase (LOR) and saccharopine dehydrogenase (SDH)^18^. Protein levels of AASS are unchanged between podocytes and tubule mitochondria (Supp. Figure 4B), but cross-link level analysis shows significant differences (**Figure 2F**). There is a subset of cross-links that show significant increase in podocyte mitochondria. Mapping of these links on structures of AASS LOR and SDH domains in XLinkDB showed that out of 4 links 1 is a homodimeric link (K728-K732), while another link (K728-K739) is possible as both intralinks and homodimeric link. K253-K70 cross-link is more probable as a homodimeric link than intralinks. Only K253-K259 is more likely to be an intralink. Taken together these cross-links indicate that in podocyte mitochondria there are more assembled multimers, which could indicate higher activity based on the reports that enzyme is only active as an oligomer^19^. The majority of studies of the role of lysine in kidney physiology and pathology use whole kidney tissue or isolated proximal tubules. Our quantitative comparisons of tubule and podocyte interactomes and proteomes could help study lysine metabolism at higher resolution in kidney. The proteins in lysine degradation are also part of tryptophan metabolism with an addition of several others (Figure 2E, Supp. Figure 4D). One of the proteins that is in the tryptophan pathway, but not in lysine degradation, is kynurine 3-mono-oxygenase (KMO). KMO is one of the few proteins differentially expressed in podocyte and tubule mitochondria with higher abundance in podocytes (**Figure 4G**). Although high expression of KMO in proximal tubules on both transcript and protein level has been reported before, our data points to higher expression of KMO per mitochondria in podocytes^20^. This could be explained by the fact that both single cell transcriptomics and kidney tissue immunohistochemistry would show higher level of KMO in tubules due to the higher overall mitochondrial content. It has been shown that loss of KMO causes proteinuria directly linked to podocytes^21^. Tryptophan degradation metabolites have also been linked to renal failure^22^. Overall, this illustrates how this dataset is uniquely positioned to distinguish podocyte specific mitochondrial biology from the otherwise high background of mitochondrial abundance imparted by tubule cells.

### Glutamate dehydrogenase activity, but not protein levels are decreased in podocytes

Glutamate dehydrogenase (DHE3), encoded by *Glud1* gene, is a mitochondrial matrix enzyme responsible for oxidative deamination of glutamate to produce alpha-ketoglutarate (aKG), an intermediate of TCA cycle (**Figure 3**.)^23^. The expression of DHE3 in mitochondria between podocytes and tubules is the same (**Figure 3**.), however, on the cross-link level there are extensive differences (**Figure 3**.). While there are cross-links that show no change similar to protein levels, others are increased indicating conformational differences. There is also a subset of cross-links that are significantly decreased in podocyte mitochondria. These cross-links are localized to the antenna regions of hexameric DHE3. We have previously shown that the same cross-links are decreased in female mouse muscle mitochondria with age^8^ and we were able to demonstrate that these cross-links increase with treatment by adenosine diphosphate (ADP), a well-known activator of DHE3 activity^24^. To investigate further DHE3 activity in podocytes and tubules we performed an activity assay which measures NADH absorbance as it being produced during the DHE3 oxidative deamination reaction, using both mitochondria from the same mice as were cross-linked, as well as 3 more pairs. As predicted by cross-link levels, time to reach half maximum of the NADH absorbance in podocytes was longer compared to tubules, indicating that DHE3 activity is lower in podocytes (**Figure 3D, Supp. Figure 5A**). DHE3 is an important mitochondrial enzyme that operates as a metabolic switch by employing intricate regulatory system that, curiously, co-evolved with the antenna^23^. To investigate a possible underlying mechanism driving increased activity we have performed activity assays and quantitative cross-linking experiments treating mouse liver (which has the highest expression of DHE3 for maximum depth of coverage) mitochondria with either DHE3 activator (ADP), inhibitor (GTP), or both (**Figure 3E, Supp. Figure 5B**). The antenna cross-links increased in response to ADP or ADP + GTP treatment but not to GTP treatment alone. There was also no response to either ADP or GTP when no substrates were added (the mitochondria were previously frozen, so it is expected that due to membrane raptures small molecules like glutamate and co-factor NAD^+^ would be lost). These results indicate that antenna cross-links are responsive to ADP activation but not to GTP inhibition, indicating that differences in DHE3 activity in podocytes in tubules might be driven by

**Figure 3.**
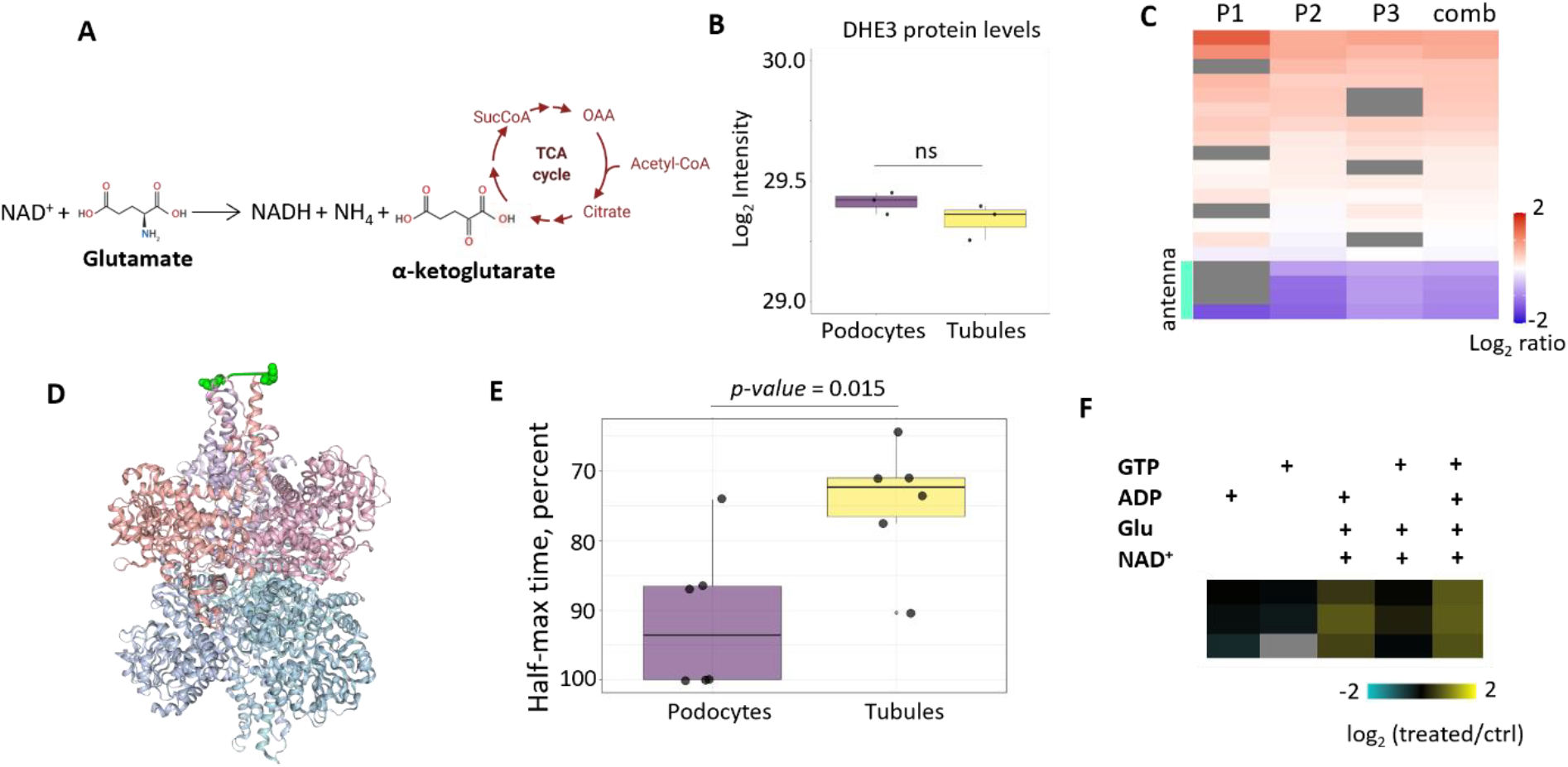
DHE3 cross-links and activity are decreased in podocytes compared to tubules. **A**. DHE3 converts glutamate and NAD^+^ to NADH, ammonia and alpha-ketoglutarate, a TCA cycle intermediate. **B**. Podocyte and tubule mitochondria have similar DHE3 levels. Significance is determined by two-sided t-test with Benjamin-Hochebrg correction (ns = non-significant). **C**. Heatmap of all confidently quantified DHE3 cross-links. **D**. Structure of DHE3 with a representative antenna link. **E**. Boxplot comparing time to reach half maximum of NADH absorbance as measured by DHE activity assay in podocytes and tubules. The times are normalized to the maximum of each experiment. Statistical significance is determined by two-sided t-test. **F**. Heatmap of antenna cross-links measured in previously frozen liver mitochondria with addition of ADP, GTP and ADP+GTP.

### Concluding remarks

Here we have demonstrated, for the first time, that iqPIR based quantitative cross-linking mass spectrometry can be used to detect reproducible differences in interactomes originating from the different cell types within the same organ. Mitochondria of kidney podocytes and tubules show few differences on a proteome level but more extensive differences at the interactome level. Using these cross-linking data we were able to identify pathways that are differentially regulated in podocyte mitochondria but that are otherwise masked in proteome data due to the overwhelming abundance of mitochondria in tubules relative to podocytes. Proving that these data can be used to make mechanistic predictions of cell-specific mitochondrial funcitons, we have discovered podocyte versus tubule mitochondria differential activity of glutamate dehydrogenase through cross-links in its antenna region and confirmed this by orthogonal assay. Data generated in this study will be publicly available and could be used by the community to gain a deeper understanding in differences between mitochondrial biology of tubules and podocytes. In the future, interactomes of other intra-organ cell types could be studied with this method to gain new insights into their biology.

## METHODS

### Animal care

All strains of mice used in this study are on a C57BL/6 background. Founding breeder mice used to produce all genotypes of mice included in this experiment were obtained from The Jackson Laboratory. The NPHS2-cre transgene was introduced into the colony with mice of the B6.Cg-Tg(NPHS2-cre)295Lbh/J strain (The Jackson Laboratory, strain #008205), the CDH16-cre transgene with mice of the B6.Cg-Tg(Cdh16-cre)91Igr/J strain (The Jackson Laboratory, strain #012237), and the MITO-HA mutation with mice of the B6.Cg-Gt(ROSA)26Sortm1(CAG-EGFP)Brsy/J strain (The Jackson Laboratory, strain #032290). Mice were housed at 20°C under diurnal conditions with ad libitum food and water in an AAALAC-accredited facility with supervision by the Institutional Animal Care and Use Committee at the University of Washington. Genotypes of all animals in this study were obtained through the automated genotyping service provided by TransnetYX.

### Cell-specific MACS HA-pulldown for Mitochondrial Isolation

Mice were sacrificed, and kidneys removed, decapsulated on a cold marble block, and weighed before being placed into ice-cold KPBS. 100 mg of tissue per column was weighed and then placed into 1 mL of KPBS in a glass Dounce homogenizer on ice and ground to a slurry. Tissue homogenates were poured through a 100 uM cell strainer to remove large debris and then mitochondria were isolated with the MACS column system (Milteny, Germany) using a proprietary kit designed specifically for mitochondrial isolation and following all instructions with two exceptions. First, to isolate HA-tagged mitochondria, the mitochondrial kit components were used, but with Milteny anti-HA beads substituted in place of the TOMM22 beads. Second, all bead incubations were performed for 45 minutes. After immunoprecipitation of mitochondria following kit protocol, mitochondria pellets are washed twice and resuspended in 1.5 mM EGTA, 3 mM MgCl2-6H2O, 10 mM KH2PO4, 20 mM HEPES, 110 mM Sucrose, 100 mM Mannitol, 60 mM K-MES, pH 7.1. For podocyte mitochondria, two columns were run to ensure enough mitochondria are collected for cross-linking. For tubule mitochondria, one column yields sufficient mitochondria. Following the modified kit protocol, time from animal euthanasia to isolated respiring mitochondria is ∼90 minutes.

### Cross-linking of mitochondria

Mitochondria were cross-linked and processed as reported before.^8^ Briefly, mitochondria were cross-linked with either RH iqPIR reagent (podocytes) or SH reagent (tubules). Reaction was allowed to proceed for 30 minutes. Mitochondria were then lysed in 8 M urea, sonicated, reduced with TCEP and alkylated with IAA. Protein content was measured with Bradford assay and then protein from podocytes and tubules were mixed pairwise in 1:1 to maximize total protein per pair (podocyte sample with highest protein amount was mixed with tubule sample with highest protein amount and so on). Mixed samples were digested overnight with trypsin and cleaned up using c18 SepPak coloumns (Waters). Peptides were then fractionated using SCX column (Luna column, Agilent HPLC) and fractions containing cross-links or deadends were used for avidin capture. Fractions were then separated on Aquity nanoHPLC (Waters) and analyzed on QE plus mass spectrometer (ThermoFischer Scientific). The data was acquired in DDA mode with top 5 peaks with charge >3 chosen for MS^2^.

### Data processing and analysis

Mass spectrometry data was then searched with Mango software to identify spectra containing cross-linked peptides.^29^ The spectra was then searched with comet engine against Mitocarta 3.0 database or whole mouse database and cross-link identifications verified with XLinkProphet.^30,31^ Cross-links and deadends were then quantified with iqPIR algorithm to produce log_2_ ratio.^7^ A combined ratio was calculated by using ions from all samples. Cross-links IDs and associated quants were then uploaded into XLinkDB for structure mapping, further analysis and visualization.^32^ XLinkDB formatted table was exported for downstream analysis in R using tidyverse environment.^33^

### DIA based protein quantitation

After cross-linked mitochondrial samples were lysed and alkylated, 20 μg of each sample was set aside for whole proteome quantitative analysis. The samples were digested overnight with trypsin, peptides were then cleaned up using c18 SepPak columns, dried down, and resuspended in 0.1% formic acid to a final concentration of 0.5 μg/μL. DIA data was acquired and analyzed as reported before.^10,11^ Briefly, a pooled sample was made by mixing all samples in equal amounts. Gas phase fractionated (GPF) narrow window (4 m/z) libraries were then acquired using the pooled sample. Wide window (24 m/z) DIA runs were then acquired for each sample. The chromatographic libraries were constructed in Encyclopedia using GPF runs searched against mouse Mitocarta 3.0 database or whole mouse database downloaded form Uniprot.^9^ Wide window runs were searched against the libraries. Table with protein quantitation was then imported into R for downstream analysis.

### Glutamate dehydrogenase assay

Mitochondria were either prepared from fresh isolations, or thawed on ice from frozen preparations and 25 μg from each sample was prepared for glutamate dehydrogenase assay according to manufacturer instructions (Millipore Sigma MAK099). The absorbance at 450 nm was then measured every two minutes for 4 hours on Cytation 5 (Biotek) in 96-well plate. The data was then imported into R for further analysis. Half-time absorbances were calculated for each sample as a time of the first measurement after half maximum of each sample’s absorbance is reached.

### Liver mitochondria treatment with ADP/GTP and cross-linking

Liver mitochondria was isolated as described in Fezza et al^34^. Briefly, tissue was homogenized in glass homogenizer, cellular debris were cleared by centrifugation at 800 g at 4C and mitochondrial pellet was isolated by centrifugation at 10000g. Mitochondria were washed in isolation buffer and stored in -80C until cross-linking experiments. For treatments and cross-linking experiments mitochondria were thawed on ice and split into pools. Substrates and inhibitors/activators were added as described in **Figure 3**.. Mitochondria were then cross-linked as described for podocyte and tubule mitochondria cross-linking. Samples were processed and analyzed in a similar fashion.

## ACKNOWLEDGEMENT AND FUNDING

This work has been supported by the following grants: K01 AG062757-01 (MTS), K01 AG073470-01A1 (MSC), R21 1R21DK128540 (MTS, MSC), P01 AG001751 (Rabinovitch), AHA 23PRE903119 (AAB). Authors appreciate thoughtful discussion with members of Bruce lab.

## SUPPLEMENTAL FIGURES

**Supplemental Figure 1.**
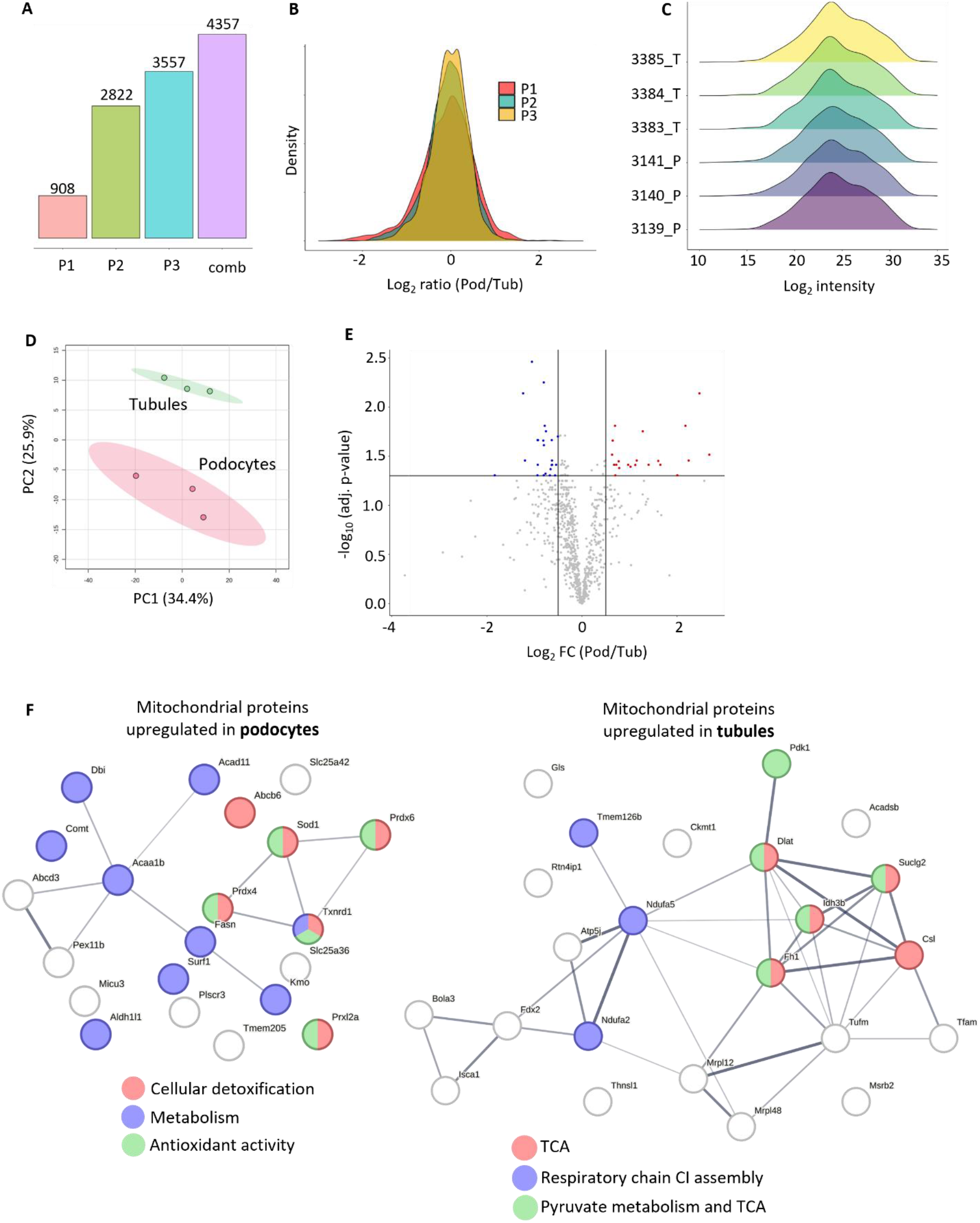
Interactome and proteome quantitation in podocyte and tubule cells searched against Mitocarta 3.0 database. **A**. Number of cross-links quantified before strict quality cut-offs (95% confidence <= 0.5, >= 4 ions used for a ratio). **B**. Distribution of log_2_ ratios in each pair using confidently quantified cross-links. **C**. Distributions of normalized log_2_ transformed intensities of protein quants for each sample (podocyte and tubule samples marked with “_P” and “_T” respectively. **D**. PCA plot shows separation of podocyte and tubule samples based on protein quants. **E**. Volcano plot of proteins significantly upregulated in podocytes (red, right) and in tubules (blue, left). Significance is determined by two-sided t-test with Benjamin-Hochberg correction (p-value = 0.05) with |log_2_ FC| >= 0.5. **F**. STRING networks of proteins upregulated in podocytes (left) or tubules (right). Some of the significant pathways are highlighted.

**Supplemental Figure 2.**
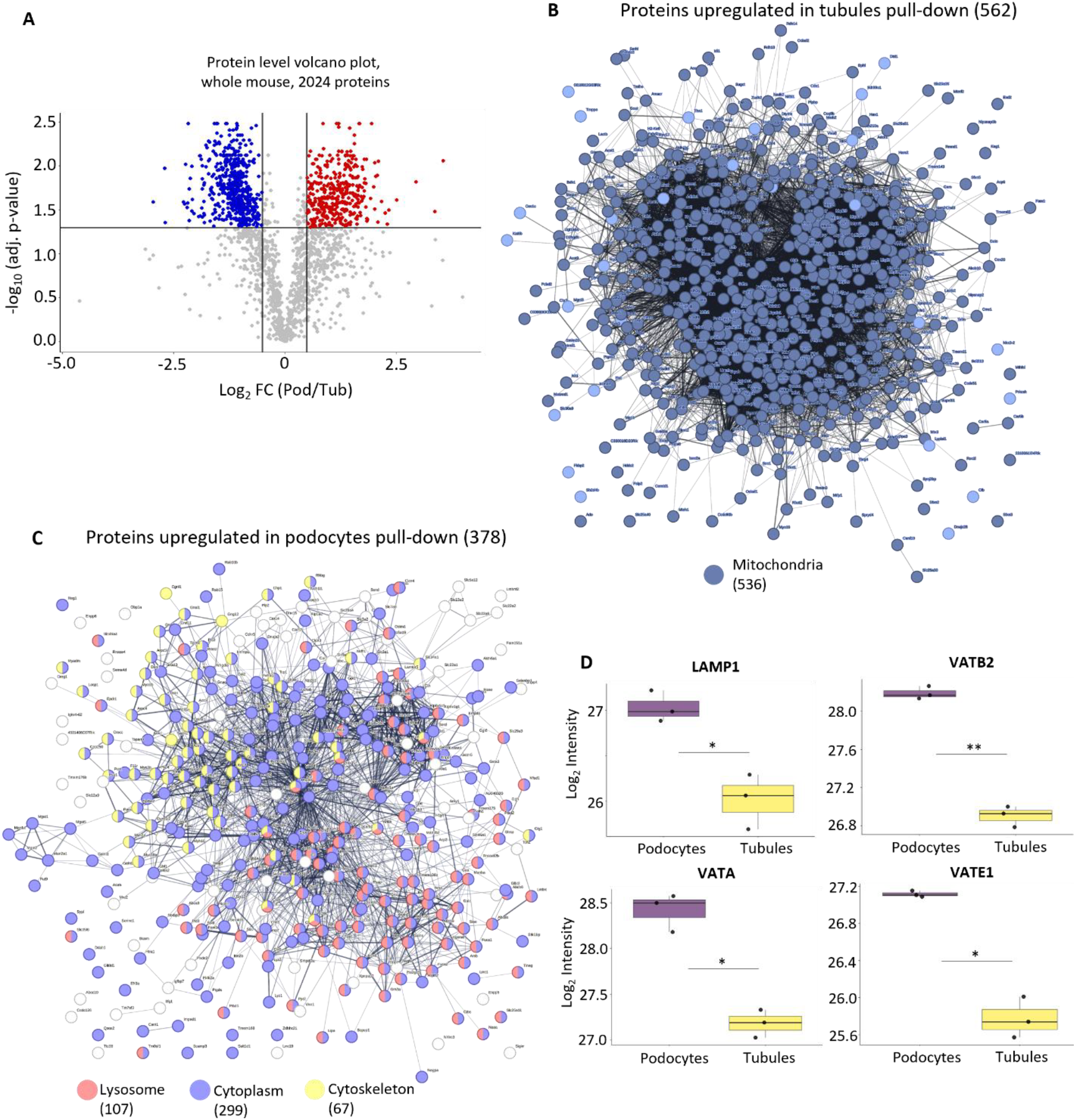
Proteomics analysis utilizing whole mouse database searches in the podocyte and tubule pull-down. **A**. Volcano plot demonstrating differential proteins between podocyte and tubule pull-down. Significance is determined by two-sided t-test with Benjamin-Hochberg correction (p-value = 0.05) with |log_2_ FC| >= 0.5. **B**. Proteins significantly increased in tubule pull-down. Most proteins are annotated as mitochondrial. **C**. Proteins significantly increased in podocyte pull-down. Lysosomal, cytosolic and cytoskeletal proteins are enriched in this pull-down. **D**. Boxplots comparing select lysosomal protein levels between podocytes and tubules. T-test determined significance is indicated (* = 0.05, ** = 0.01).

**Supplemental Figure 3.**
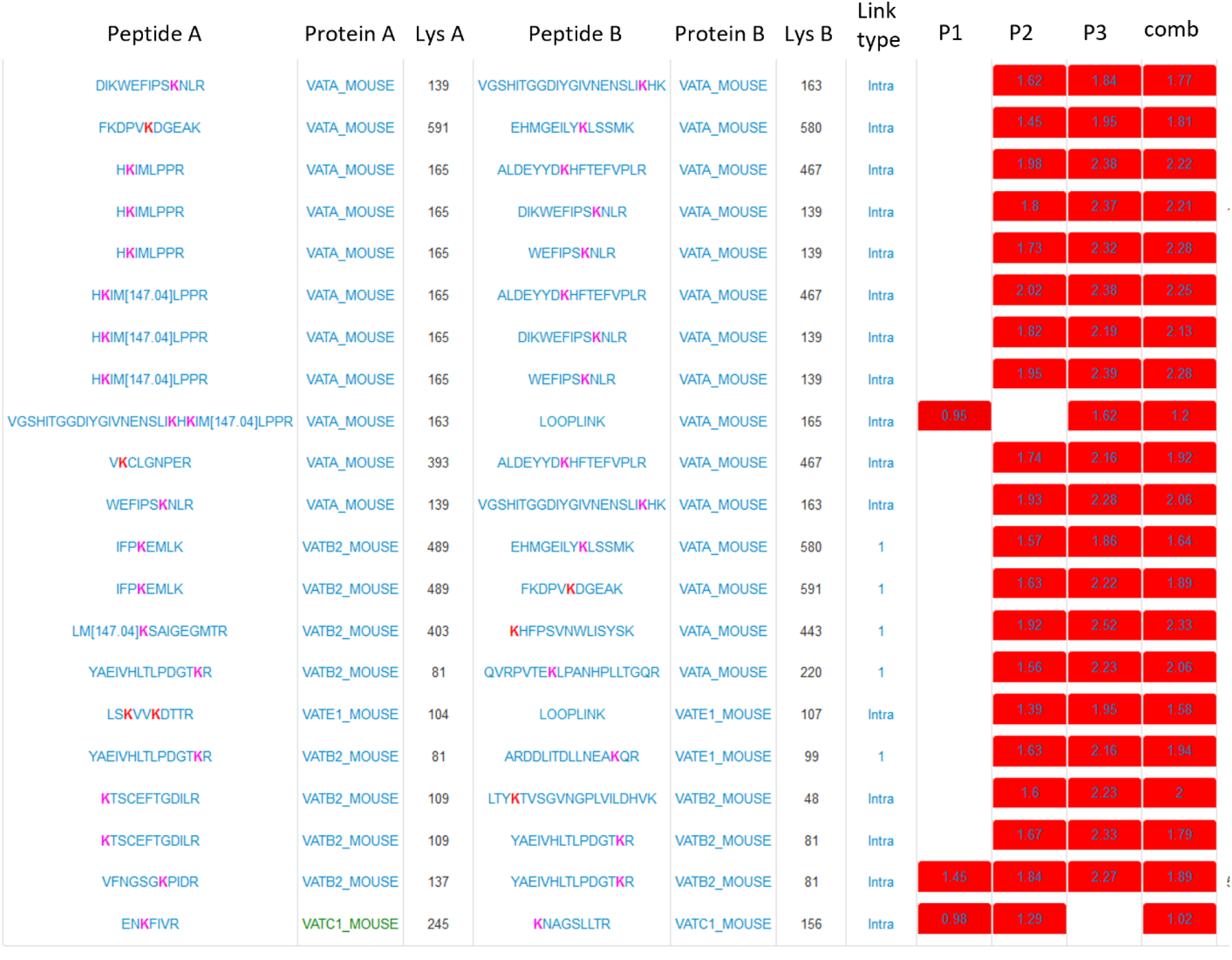
Cross-links from lysosomal proton pump show increase in both intra and intersubunit links in podocytes. Link type “1” indicates inter-subunit cross-links.

**Supplemental Figure 4.**
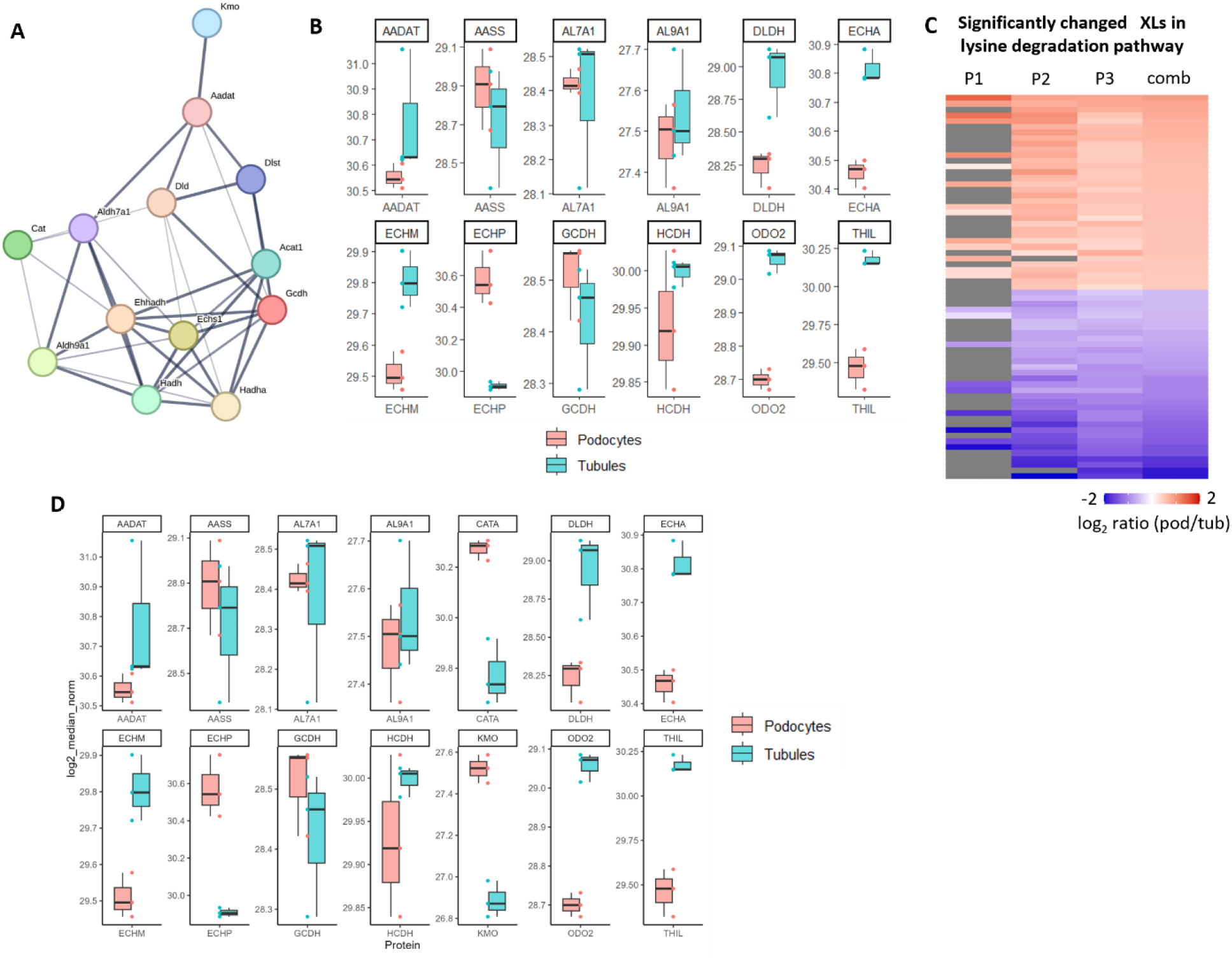
Lysine and tryptophan metabolism pathways are altered between podocyte and tubule mitochondria. **A**. STRING network of proteins involved in lysine metabolism KEGG pathway with differential cross-links **B**. Proteins from lysine degradation pathway with differential cross-links. **C**. Heatmap of differential cross-links in lysine degradation pathway. **D**. Protein level comparison of tryptophan metabolism proteins that had differential cross-links.

**Supplemental Figure 5.**
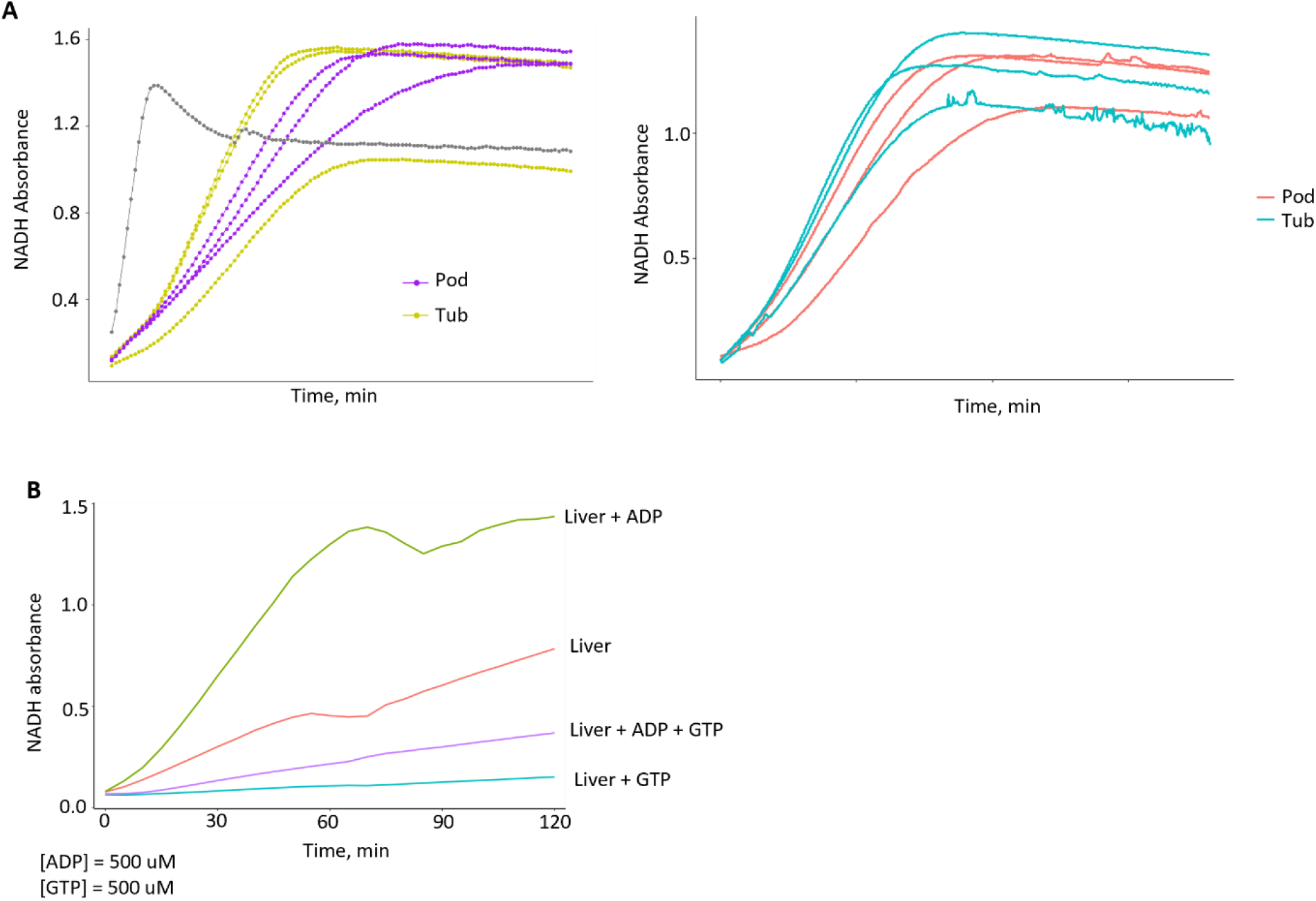
Assay to measure DHE3 activity. **A**. NADH absorbance measured at 450 nm in podocytes and tubules in the mitochondria used for cross-linking experiments (left) and additional samples (right). **B**. NADH absorbance measured in liver mitochondria after pre-incubation with substrates and addition of DHE3 inhibitor (GTP) and activator (ADP) or a combination of both.

